# Survival analysis of pathway activity as a prognostic determinant in breast cancer

**DOI:** 10.1101/2021.05.19.444688

**Authors:** Gustavo S. Jeuken, Nicholas P. Tobin, Lukas Käll

## Abstract

Most activities in a cell are dependent on sets of biomolecules, often referred to as pathways. Pathways are useful to study as their activities are frequently more directly related to the behavior of the cell than their components. Traditional pathway analysis gives significance to differences in the pathway components’ concentrations between sample groups. Here we instead advocate a singular value decomposition-based method for estimating individual samples’ pathway activities that allow us to investigate more advanced statistical models. Specifically, we investigate the pathway activities’ association with patients’ survival time based on the transcription profiles of the METABRIC dataset. Our implementation shows that pathway activities are better prognostic markers for survival time in METABRIC than the individual transcripts. We also demonstrate that we can regress out the effect of individual pathways on other pathways, which allows us to estimate the other pathways’ residual pathway activity on survival.

## Introduction

Modern high throughput techniques in molecular biology enable the measurement of thousands or even millions of analytes in any sample of biological origin. Such wealth of data makes it possible to describe samples very accurately, with a precision that opens the door to new understandings of the mechanisms governing biological and medical processes.

Although this abundance of data indicates a possibility for acquiring knowledge, the sheer number of dimensions in such measurements also presents us with many challenges: when analyzing high-dimensional vectors with statistical methods one easily faces the curse of dimensionality, i.e. that the sample space grows exponentially with each added dimension [1]. This becomes a problem when the number of samples is lower than the number of measured analytes. Further, in e.g. differential expression analysis, which tests the concentration differences of each measurement separately, we to some degree run into decreased sensitivity due to multiple hypothesis testing.

One of the more promising ways of alleviating these problems is through pathway analysis [2, 3]. Proteins frequently operate in an orchestrated manner and phenotypes are frequently the consequence of sets of proteins and not just of single-proteins. Metabolic pathways, or other aggregated biological knowledge that groups analytes, provide a model-driven way of combining molecular information and thus also the measurements from a high throughput experiment in a biologically meaningful way.

Traditional pathway analysis first determines the quantitative differences in gene expression between patient groups, and subsequently either tests the significantly differential genes annotation for enrichment in the tested pathway [7, 8, 9], or tests if the genes belonging to a pathway have more extreme differences than other genes, using so-called gene-set enrichment analysis [3]. Both types of analysis leave the user with a significance value for the analytes in a pathway annotation being differentially abundant under the different conditions. Yet this feels unsatisfying, as embedding the pathway analysis in the statistical test significantly limits the types of statistical tests that can be applied in the analysis.

An alternative is offered by single sample pathway analysis, which promises activity scores for each sample and pathway. The method ASSESS [10] fits two mixture models to data to quantify pathways, however, the supervised learning step of the procedure makes further statistical testing challenging. Another critique is that most pathway analysis methods derive statistical significance by comparing the behavior of genes belonging to a pathway with those that do not. Goeman & Bühlmann [11] argue in favor of self-contained tests, in which the significance of a pathway relates only to the expression of said pathway’s genes, resulting in a more restrictive null hypothesis that leads to a higher statistical power. GSVA [12] uses a competitive test for gene set enrichment. The ssGSEA [13] method, by utilizing internal ranks of expression as a basis for enrichment, is thus also not self-contained. While this is also true for singscore [14], it has a large advantage in that it does not need other samples as a background when providing each sample’s score.

Two methods, PLAGE [15] and Pathifier [16], will produce self-contained unsupervised metrics of pathway activity. The former does so through Singular Value Decomposition (SVD) in a space formed only by a pathway’s genes. The latter takes the same approach but uses a Principal Curve instead, and while this enables non-linear gene interactions to be captured, it introduces the need for annotated baseline samples, as well as a larger dataset.

We strongly believe in the merits of doing statistical analysis on a pathway level, as the function of a pathway is often much better understood than that of individual genes, and statistical operations on a pathway-level are often more directly related to the biology of the problem. Cell proliferation, as an example, has been established as one of the hallmarks of cancer [4, 5]. It is a complex, systemic process that involves many different mechanisms, in different parts of the cell, as well as signaling pathways that regulate them. In breast cancers, it has been shown that one can separate Luminal A from Luminal B subtypes based on the level of proliferation [6]. Thus, when investigating how molecular profiles of tumors affect patients, one may want to look at proliferation as a whole, instead of focusing on individual analytes.

The availability of methods that generate sample-specific pathway scores opens the way for many statistical methods that were previously inaccessible to pathways, yet we feel this is still an underutilized resource. We specifically think that SVD-based methods, due to the properties described above, as well as their ability to directly capitalize on gene covariation, are a strong base for further statistical analysis. Thus, we here reimplemented PLAGE’s SVD-methodology [15] for pathway summarization and studied its performance when applied to an advanced statistical method, namely survival analysis. We highlight the advantages and flexibility that working on a pathway level provides. Particularly we demonstrate how to counter the confounding effects on the pathway analysis of increased cell proliferation in severe breast cancers. We also demonstrate how one can use sunburst plots for exploratory visualization of the significance of pathway activities while maintaining a pathway hierarchy.

## Materials and methods

### METABRIC Transcription profiles, and their annotation

Normalized gene expression and clinical annotation in the METABRIC dataset were downloaded from the European Genome-phenome Archive. This data consists of microarray reads from 1992 breast cancer specimens, primarily fresh frozen, together with the clinical annotation, including survival information, of the respective patients. Twelve samples were reported twice in the dataset [17] and these were removed from our analysis, but otherwise, the entire cohort was used.

Pathway annotations were retrieved from the Reactome database [18], version 76, with annotations as Ensembl gene IDs. The pathways’ gene ID annotations were converted to Illumina probe IDs (HT_12_v4), through BioMart [19], by assuming that any transcript associated with each Reactome protein’s underlying gene was associated with the pathway.

The clinical endpoint for this study was breast cancer-specific survival (BCSS) defined as patients who have not died from breast cancer in the study period from the date of surgery to the end of follow-up.

### Estimation of Pathway Activities

We followed the PLAGE [15] method’s singular vector decomposition strategy. Let 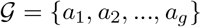 be the set of all measurements we want to study, the Illumina probes in our case, and let 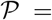 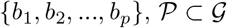 be the subset of measurements that are associated with our pathway of interest. We define **A**_*g×m*_ as the matrix of log-transformed and standardized measurements of our samples (where *m* is the number of samples), and **B**_*p×m*_ as a matrix constructed by using only the rows in **X** that are present in 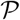. We then decompose **B** using a Truncated Singular Value Decomposition (SVD):

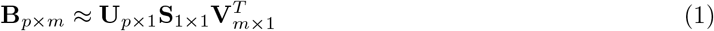

Note that there is a slight difference in preprocessing compared to Tomfohr *et al*.[15] which use a **A**_*g×m*_ that contains standardized measurements that were not log-transformed. Here we name the pathway’s left singular vector **U** as an eigensample, and the right singular vector **V** as an eigengene, following the nomenclature in Wall *et al*. [20]. Note that the eigengene contains one vector element per sample, and we will use these vector elements as a measure of pathway activity. Practically we here make use of the *scikit-learn* python package [21].

### Proportional Hazards Model

Cox’s proportional hazards model [22, 23] relates the survival function *S_i_*(*t*) of patient *i* to the value of any of its covariates *X_i_* through the following:

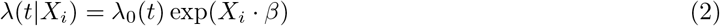

Where *λ*(*t*) is the hazard function and is defined as the rate of mortality at time *t*, for patients that have survived up to that time, and *λ*_0_(*t*) is the baseline hazard defined as *λ*(*t*|0). The hazard function is then associated with the survival function by *λ*(*t*) = *S*′(*t*)/*S*(*t*)

Here we used the pathway activities derived previously as the exploratory variables *X* for a Cox regression, doing so one pathway at a time, to study the connection between each of the samples’ pathway activities and the survival of patients. This regression gave us both a coefficient *β* showing the magnitude of the effect of a pathway, and a *p* value representing the statistical significance of its coefficient. The *p* values were subsequently corrected for multiple testing into *q* values [24]. In here we make use of the *lifelines* python package [25].

### Concordance Index

The concordance index (or C-index) is a generalization of the area under the curve (AUC) classifier performance that can take into account censored data. It represents the model’s accuracy in ranking the survival times of the samples[26]. It can be calculated as:

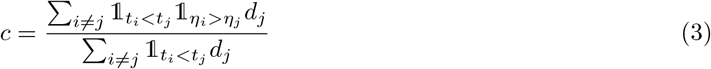

Here, for each patient *i*, we have the observed survival time, *t_i_*, and the censoring variable, *d_i_*, that takes a value of either 1 if the event of death has been observed and 0 otherwise. The indicator variable 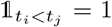 if *t_i_* < *t_j_* and 0 otherwise. The variable *η* is the hazard score for each sample, calculated as *η_i_* = *X_i_* · *β*. The C-index is calculated using 5-fold cross-validation: for each step, the coefficients *β* are fitted to 80% of the data, the hazard score *η* is calculated for the samples in the holdout data and the index is obtained by comparing it to their survival status. Just as for AUC, a concordance index of 0.5 corresponds to a null prediction and 1 to a perfect prediction.

## Results

### A pathway-level survival analysis

We implemented a method for evaluating a pathway activity’s influence on survival. The method is generic for any censored data, however, here we demonstrate its efficiency on the METABRIC breast cancer dataset. The transcription profiles of the 1980 breast cancers in the METABRIC data were downloaded and grouped according to the Reactome database into pathway groups. After the operation, each pathway group contained the transcripts corresponding to proteins in each pathway, and as there are overlaps between pathways in Reactome, this means that each transcript could appear in multiple pathways. The expression matrix of each pathway was factorized into eigensamples (left-singular vectors) and eigengenes (right-singular vectors) using the first eigenvector of a Singular Value Decomposition (SVD) following Wall *et al*. [20]. The eigensample represents the linear combination of genes and the eigengene the linear combination of samples that best explains the variance of the expression matrix of the pathway. This slightly backward naming convention stems from the idea that an eigengene models the variation of samples of a typical gene, while an eigensample models the variation of the genes in a typical sample.

We then project each sample into this eigengene and use this result as a representation of the sample’s pathway activity. These pathway activities together with the survival information for each patient were fed into a Cox model for proportional hazards, which regresses these values against the survival information (survival time and disease-specific death) to obtain both the regression coefficient and the statistical significance for the effect of each pathway activity on patient survival. The significance, first obtained as *p* values were multiple testing corrected into *q* values [24]. Fig 1 gives an overview of the procedure.

**Figure 1:**
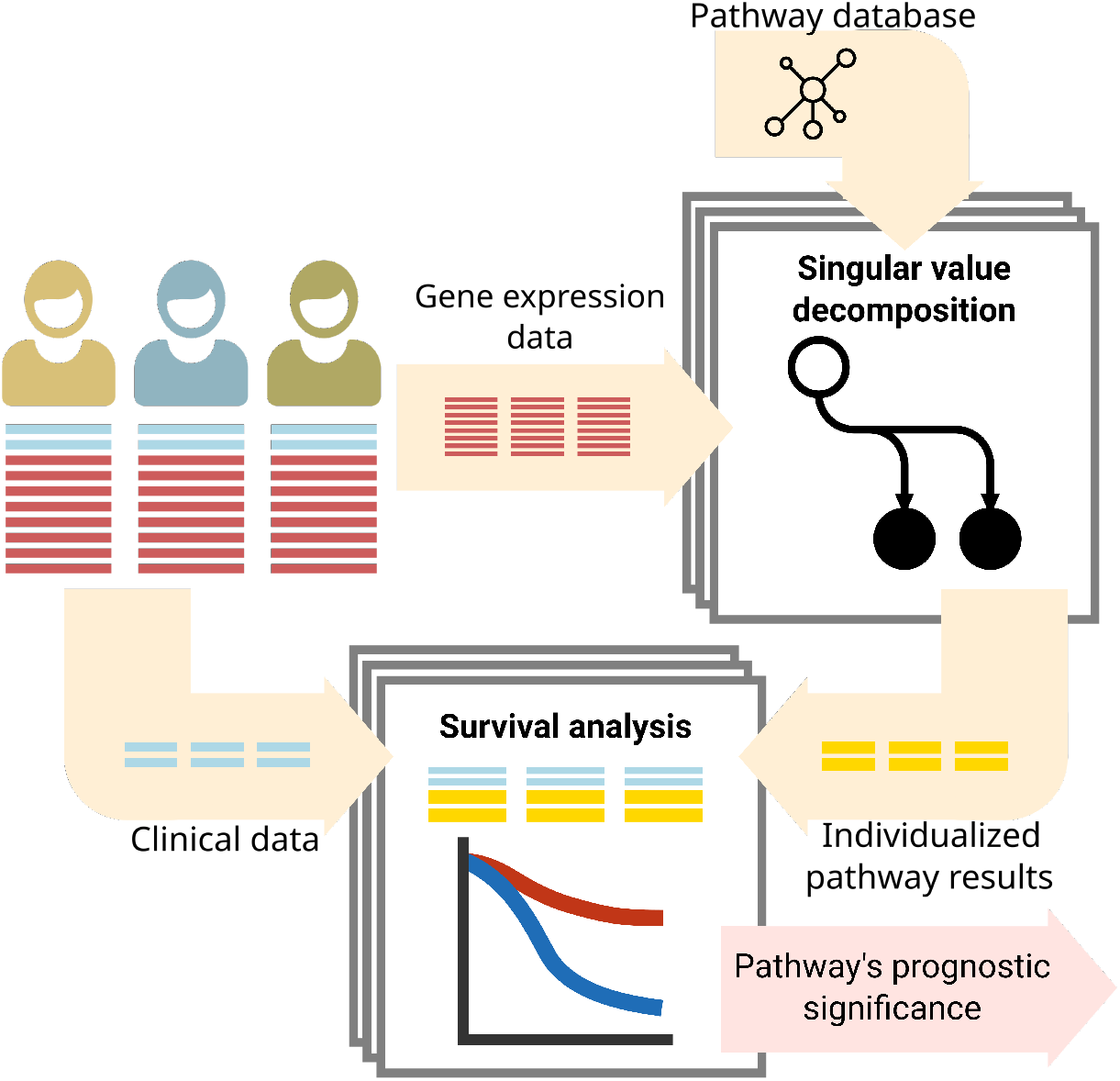
A method to analyze the coupling of pathway activity on patient survival. We used singular value decomposition to give, for each sample and pathway, an individualized pathway readout. We then combine those readouts with the survival information to perform survival analysis on a pathway level.

Before we investigate the output of our model, we would like to motivate the choice of SVD as a means to capture pathway activity. As an example, let’s consider the pathway ‘Metabolism of folate and pterines” consisting of 26 proteins. When investigating the covariation matrix of the transcripts and the survival time for 907 samples from patients that were deceased in the METABRIC cohort, we found both examples of transcript levels that correlate positively and negatively with the survival time (Fig 2). By having the probe reads ranked by their contribution to the eigensample, we see that the eigensample captures information from genes that have both a positive and negative influence on survival. We also see that probes that have a negative contribution to the eigensample also have a negative influence on survival, while the opposite is also true. It is the linear combination of these expressions, obtained in an unsupervised manner, that is used for later survival analysis.

**Figure 2:**
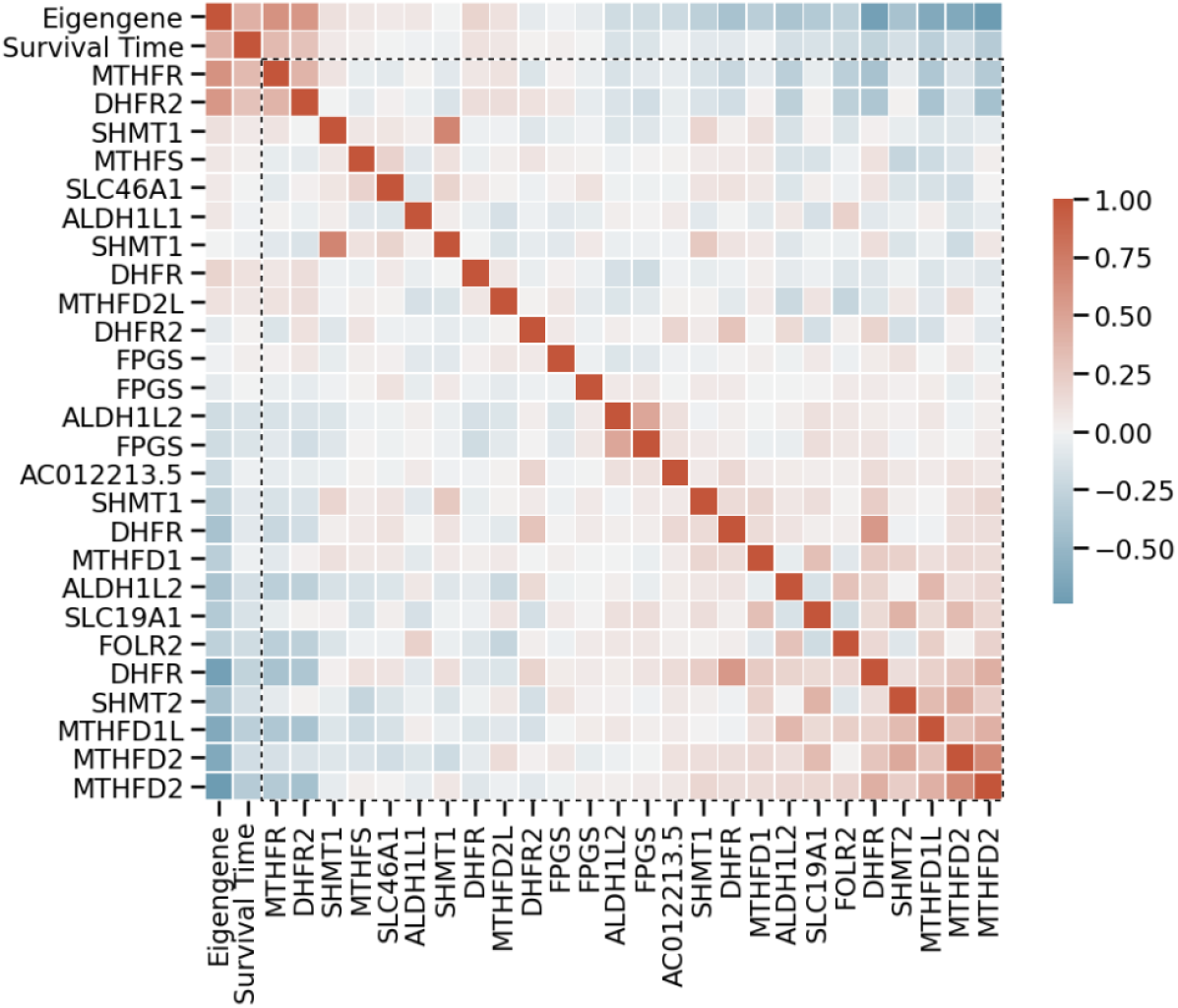
Heatmap of the coefficients of the Pearson correlation matrix between the variables survival time, the eigengene of the gene expression values, and the gene expression values within the pathway ‘Metabolism of folate and pterines” for samples from diseased patients. We sorted the genes in the expression matrix by their contribution to the eigensample and encompassed them with a dotted line. The patients’ survival time both correlates and anti-correlates with the different genes. However, the eigengene captures the covariational trend within the expression data and correlates well with the patients’ survival time.

### Analysis of breast cancer data set

The transcription profiles of the 1980 breast cancers in the METABRIC data were downloaded, and they were analyzed with our method. Supplementary Table S1 list the pathways and their prognostic significance, and the results are also available as an interactive plot (https://statisticalbiotechnology.github.io/metabric-pathway-survival/results.html). We found 1030 pathways (out of the 2214 pathways in Reactome) that were associated with patient survival with *q* ≤ 0.05. We note that most pathways involved with the Cell Cycle are indeed associated with survival, as one would expect, as well as pathways relating to DNA replication and DNA repair.

This result is unsatisfying, as we already know that Cell proliferation is a driver of cancer [4], and it specifically has a strong influence on survival. We also know that it affects most other aspects of cancer cell activity, making it hard to disentangle this signal from other important processes that affect the survival of patients. Luckily we have scored the activity of all pathways and can use them to highlight associations that otherwise are drowned by the more trivial background of cell proliferation.

Traditionally proliferation is quantified by the transcription of marker genes, like MKI67 [27]. However, here we instead used the pathway activity of the “Diseases of the mitotic cell cycle”, calculated by our model, as a proxy for abnormal cell proliferation. We then regressed out the influence of proliferation on other pathway activities, by adding an independent variable for cell proliferation in our Cox’s regression model for each pathway.

After regressing out our measure of proliferation the number of significant (*q* ≤ 0.05) pathways, as expected, is reduced to 264 (Supplementary Table S2, (https://statisticalbiotechnology.github.io/metabric-pathway-survival/results_proliferation.html). We see that the operation removes the significance of most pathways relating to the cell cycle and DNA replication. Also, several other pathways’ significance diminishes, e.g. the “Metabolism of folate and pterines” pathway became insignificant, indicating that its prognostic power was mostly driven by cell proliferation (when removing that confounding effect of cell proliferation, its effect on the survival of the patients is removed, from *q* = 9 · 10^−13^ to *q* = 0.17). However, pathways relating to cell pH regulation due to respiratory oxidation [28] (e.g. “Bicarbonate transporters”, from *q* = 2 · 10^−10^ to *q* = 7 · 10^−5^) and HER2-signaling [29] (e.g. “GRB7 events in ERBB2 signaling”, from *q* = 3 · 10^−8^ to *q* = 7 · 10^−5^) remain highly significant after the operation. This demonstrates an advantage of calculating individual measurements of pathway activity for each sample, as we can subtract the effects of known confounders.

### Pathway’s eigensamples appear stable and their estimates appear well calibrated

For the results to be meaningful, the eigensample should be stable, i.e. if the decomposition is repeated for similar samples, a similar eigensample should result. This is especially important for datasets that are smaller than the ones used here. To simulate this, we randomly subsampled 20% (396 samples) of the tumors 100 times, each time performing the same decomposition, and made pairwise comparisons of the resulting eigensamples using the cosine distance (Supplementary Fig S1). We found that most eigensamples were indeed stable, and most importantly, the most significant pathways in respect to survival prognosis were all stable (Supplementary Fig S2).

We also performed a permutation test to check the calibration of the *p* values coming from the Cox regression model. We randomly selected 100 pathways by their regression *p* value, by first sampling *a* uniformly in the range [−5, 0] and selecting the pathway with the *p* value closest to 10^−*a*^, then for each pathway the association between the gene expression values and the survival information was permuted 100/*n* times, where *n* is the regression’s *p* value. The fraction of permutations with a more extreme outcome was then compared to the original Cox model’s *p* value. The results (Supplementary Fig S3) indicate that our model is indeed well-calibrated for the null hypothesis that there is no association between gene expression in the form of the pathway activation and patient survival.

Finally, to test whether the signal we see comes from a meaningful set of transcripts, as opposed to any set of transcripts, we performed a gene set permutation analysis (Supplementary Note S1), which confirmed the relevance of the manually curated sets.

### Pathways are more predictive of survival than individual transcripts

We also tested how well eigengenes can be used to predict the survival of each patient. We do this by, for each pathway, performing 5-fold cross-validation using the Cox model to predict survival on the holdout data, measuring the success of the prediction using the Concordance Index. Fig 3A shows the distribution of the Concordance Index after performing the cross-validation on all pathways and contrasts it to the distribution of the concordance index obtained by doing the same process with individual transcripts. We see that there is a general gain in predictive power when combining the information contained in individual transcripts using our method. When we compare pathways against only their constituent transcripts, we note that the presence of a strongly predictive transcript is a necessary but not sufficient condition for a highly predictive pathway, e.g. the top 10 best predictive pathways contain 6 of the top 10 transcripts, yet these same transcripts are present in pathways with no predictive power (0.49 < *CI* < 0.51).

**Figure 3:**
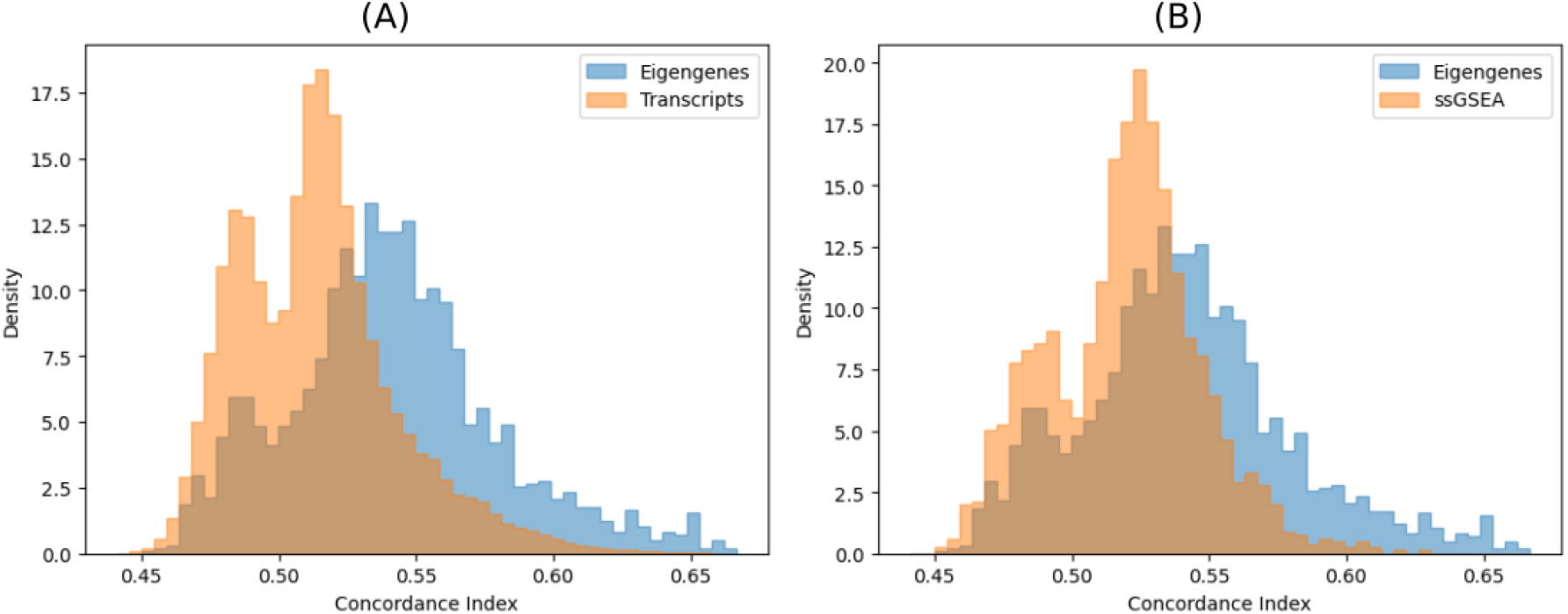
Comparison of how well eigengenes predict survival. A histogram of the concordance indices was calculated with 5-fold cross-validation of Cox regressions based on eigengenes compared to (A) individual transcripts and (B) ssGSEA enrichment scores.

### Eigengenes are more predictive of survival than enrichment scores

We compared our method to the alternative method of using the enrichment scores of single-sample geneset enrichment analysis (ssGSEA)[13] instead of eigengenes as an independent variable in a Cox-regression. To obtain the enrichment scores, we used ssGSEA as implemented in https://github.com/broadinstitute/ssGSEA2.0 with its default parameters. We found our eigengenes more sensitive than ssGSA, with 1030 vs 352 reported pathways with *q* values below 5%. We also found that the concordance indices of our model outperformed the ones produced by the ssGSEA-fed model (Fig 3B).

### An interactive visualization of pathway-level results

We know that biological pathways, as described by pathway databases, are not independent, i.e. genes can belong to multiple pathways at the same time. Moreover, the relationship between pathways also has a hierarchical nature, and in many pathway databases, large pathways are composed of smaller, connected pathways. This information is lost when we present a table with the results of our analysis, and for this reason, we developed an interactive visualization of the results based on a sunburst chart that highlights not only the numerical results but also the hierarchy and relationship between pathways. Fig 4 shows a static view of our interactive figure, the full plot is available at https://statisticalbiotechnology.github.io/metabric-pathway-survival/

**Figure 4:**
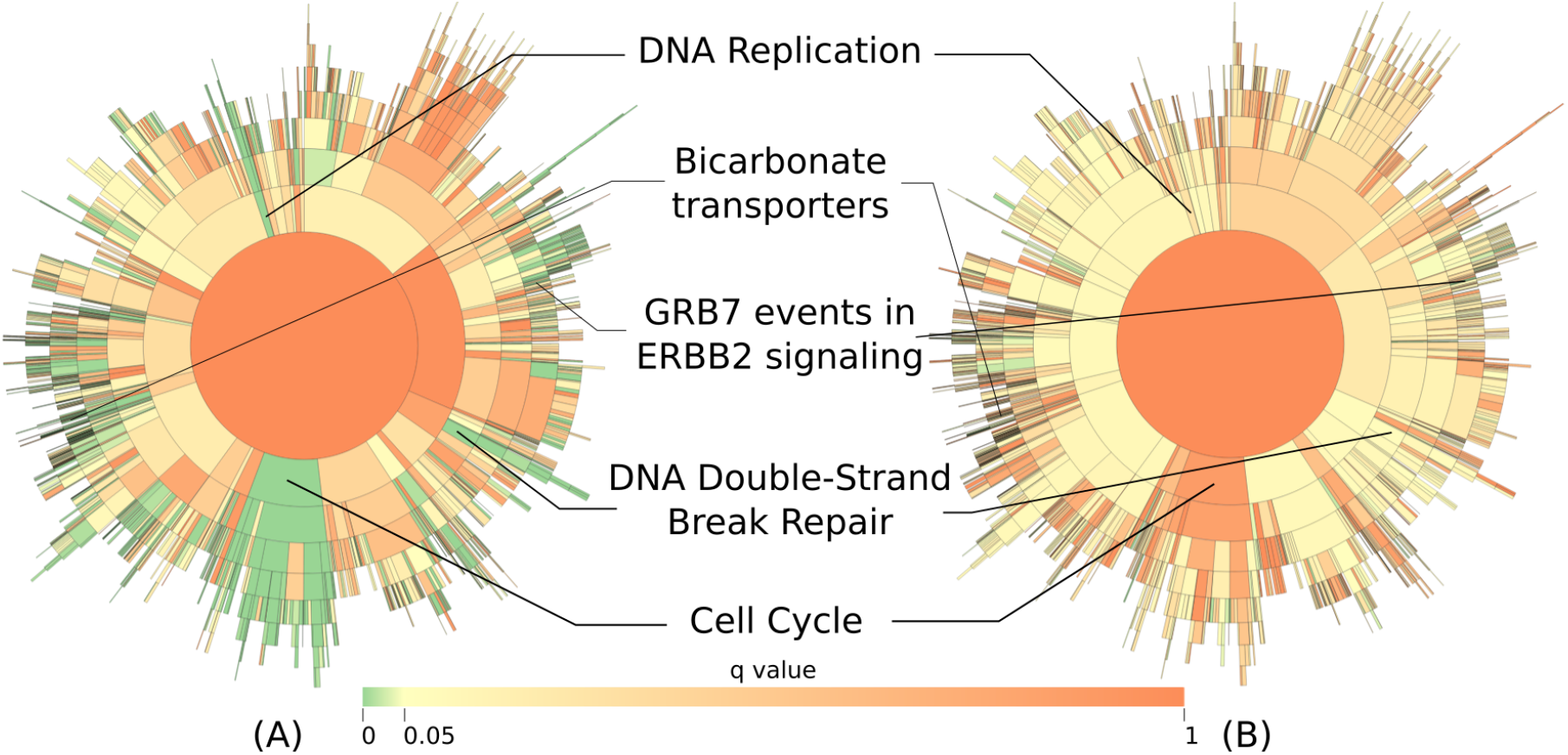
Sunburst diagrams offer an exploratory visualization of the influence of the pathways in the Reactome hierarchy on the survival of patients. We use an interactive sunburst chart to visualize and explore the result. This allows us to navigate the pathway hierarchy and see the relationships in a more natural way than e.g. in a static table. The sunburst (A) without and (B) with the pathway “Diseases of mitotic cell cycle” regressed out. Interactive versions of the same figures are available from https://statisticalbiotechnology.github.io/metabric-pathway-survival.

## Discussion

Here we have reimplemented the PLAGE method for deriving a quantitative value of pathway activity for each sample, and we demonstrated its usefulness when associating pathway activity to patient survival. This is done by singular value decomposition of the expression of genes inside a pathway, using the resulting eigensample to ascribe a pathway activity to each sample, and testing its association to the survival of patients using a Cox Proportional Hazards model.

Over the years databases of metabolomic pathways have been collected, condensing knowledge of molecular biology into causative relations. We here have made use of such relations, or at least their ability to group molecules in functional relevant subsets. Our idea is that the correlation of the concentration differences of such a subset of gene products can be seen as a manifestation of the same biological phenomenon. Thus, we argue that pathway-level statistics give a clearer view of how biological processes govern clinical outcomes. Another welcome property of pathway-level statistics is that we test fewer hypotheses than at the gene level, which increases the statistical power of the method.

One drawback of pathway analysis, however, is that, while gene expression is itself not independent, pathways by their own nature overlap and so are intrinsically dependent entities, making true multiple hypotheses corrections an even less trivial task. One should hence make sure to also check the significances of the sub- and super-pathways of the examined pathway in the studied pathway database. We believe that the sunburst plots presented in this study are very powerful for such considerations.

The results of our analysis showed many pathways that are involved with cell proliferation as the most significantly associated with the survival of the patients. Although this is an obvious result, it is also a welcome find that validates our method. Furthermore, our strategy to extract individual measurements of pathway activity for each sample allows us to elaborate on the statistical model of survival. By regressing out the effects of proliferation, the confounding effect of this major perturbation of our samples was removed, allowing us to see other features of our data more clearly [30].

At the heart of the method is the unsupervised dimensionality reduction that produces a pathway activation score for each sample. While we show some advantages of this idea on survival analysis, these should not be restricted to this field alone, and are suitable for application on any analysis where readouts are used to derive statistical significance.

Finally, SVD-based pathway analysis can be extended to include different modalities of data such as proteomics or metabolomics. As long as the novel data can be grouped into pathways, and the assumption of covariation within and between the different modalities hold, they can be included in the same factor analysis step and their information used for integrating pathway activity in an even broader sense.

## Acknowledgements

This study makes use of data generated by the Molecular Taxonomy of Breast Cancer International Consortium, which was funded by Cancer Research UK and the British Columbia Cancer Agency Branch. The computations were enabled by resources provided by the Swedish National Infrastructure for Computing (SNIC) at HPC2N partially funded by the Swedish Research Council through grant agreement no. 2018-05973.

## Supporting information

**S1 Appendix. A null model based on set permutation.**

**S1 Fig. Test of stability of pathways eigensamples.**

**S2 Fig. Investigation of the statistical calibration of** *p* **values from the Cox regression model**. Add descriptive text after the title of the item (optional).

**S1 Table. Significance of Cox-regression of Reactome’s pathways**

**S2 Table. Significance of Cox-regression of Reactome’s pathways, when regressing out Diseases of the Mitotic Cell Cycle pathway**

## References

[1] Altman N, Krzywinski M. The curse (s) of dimensionality. Nat Methods. 2018;15(6):399–400.

[2] Khatri P, Sirota M, Butte AJ. Ten years of pathway analysis: current approaches and outstanding challenges. PLoS Comput Biol. 2012;8(2):e1002375.

[3] Subramanian A, Tamayo P, Mootha VK, Mukherjee S, Ebert BL, Gillette MA, et al. Gene set enrichment analysis: a knowledge-based approach for interpreting genome-wide expression profiles. Proceedings of the National Academy of Sciences. 2005;102(43):15545–15550.

[4] Hanahan D, Weinberg RA. The hallmarks of cancer. Cell. 2000;100(1):57–70.

[5] Hanahan D, Weinberg RA. Hallmarks of cancer: the next generation. Cell. 2011;144(5):646–674.

[6] Prat A, Perou CM. Deconstructing the molecular portraits of breast cancer. Molecular Oncology. 2011;5(1):5–23.

[7] Tavazoie S, Hughes JD, Campbell MJ, Cho RJ, Church GM. Systematic determination of genetic network architecture. Nature Genetics. 1999;22(3):281.

[8] Dennis G, Sherman BT, Hosack DA, Yang J, Gao W, Lane HC, et al. DAVID: database for annotation, visualization, and integrated discovery. Genome Biology. 2003;4(9):R60.

[9] Griss J, Viteri G, Sidiropoulos K, Nguyen V, Fabregat A, Hermjakob H. ReactomeGSA-Efficient Multi-Omics Comparative Pathway Analysis. Molecular & Cellular Proteomics. 2020;19(12):2115–2125.

[10] Edelman E, Porrello A, Guinney J, Balakumaran B, Bild A, Febbo PG, et al. Analysis of sample set enrichment scores: assaying the enrichment of sets of genes for individual samples in genome-wide expression profiles. Bioinformatics. 2006;22(14):e108–e116.

[11] Goeman JJ, Bühlmann P. Analyzing gene expression data in terms of gene sets: methodological issues. Bioinformatics. 2007;23(8):980–987.

[12] Hänzelmann S, Castelo R, Guinney J. GSVA: gene set variation analysis for microarray and RNA-seq data. BMC Bioinformatics. 2013;14(1):1–15.

[13] Barbie DA, Tamayo P, Boehm JS, Kim SY, Moody SE, Dunn IF, et al. Systematic RNA interference reveals that oncogenic KRAS-driven cancers require TBK1. Nature. 2009;462(7269):108–112.

[14] Foroutan M, Bhuva DD, Lyu R, Horan K, Cursons J, Davis MJ. Single sample scoring of molecular phenotypes. BMC Bioinformatics. 2018;19(1):1–10.

[15] Tomfohr J, Jun L, Kepler TB. Pathway level analysis of gene expression using singular value decomposition. BMC Bioinformatics. 2005;6(1):1–11.

[16] Drier Y, Sheffer M, Domany E. Pathway-based personalized analysis of cancer. Proceedings of the National Academy of Sciences. 2013;110(16):6388–6393.

[17] Curtis C, Shah SP, Chin SF, Turashvili G, Rueda OM, Dunning MJ, et al. The genomic and transcriptomic architecture of 2,000 breast tumours reveals novel subgroups. Nature. 2012;486(7403):346.

[18] Joshi-Tope G, Gillespie M, Vastrik I, D’Eustachio P, Schmidt E, de Bono B, et al. Reactome: a knowledgebase of biological pathways. Nucleic Acids Research. 2005;33(suppl 1):D428–D432.

[19] Smedley D, Haider S, Durinck S, Pandini L, Provero P, Allen J, et al. The BioMart community portal: an innovative alternative to large, centralized data repositories. Nucleic Acids Research. 2015;43(W1):W589–W598.

[20] Wall ME, Rechtsteiner A, Rocha LM. Singular value decomposition and principal component analysis. In: A practical approach to microarray data analysis. Springer; 2003. p. 91–109.

[21] Pedregosa F, Varoquaux G, Gramfort A, Michel V, Thirion B, Grisel O, et al. Scikit-learn: Machine Learning in Python. Journal of Machine Learning Research. 2011;12:2825–2830.

[22] Cox DR. Regression models and life-tables (with discussion). JR Stat Soc Ser B. 1972;34:187–220.

[23] Kleinbaum DG, Klein M. Survival analysis. Springer; 2010.

[24] Storey JD. The positive false discovery rate: a Bayesian interpretation and the q -value. Ann Statist. 2003;31(6):2013–2035. doi:10.1214/aos/1074290335.

[25] Davidson-Pilon C, Kalderstam J, Jacobson N, Reed S, Kuhn B, Zivich P, et al.. CamDavidson-Pilon/lifelines: v0.25.8; 2021. Available from: https://doi.org/10.5281/zenodo.4457577.

[26] Uno H, Cai T, Pencina MJ, D’Agostino RB, Wei LJ. On the C-statistics for evaluating overall adequacy of risk prediction procedures with censored survival data. Statistics in Medicine. 2011;30(10):1105–1117.

[27] Parker JS, Mullins M, Cheang MC, Leung S, Voduc D, Vickery T, et al. Supervised risk predictor of breast cancer based on intrinsic subtypes. Journal of Clinical Oncology. 2009;27(8):1160.

[28] Gorbatenko A, Olesen CW, Boedtkjer E, Pedersen SF. Regulation and roles of bicarbonate transport in cancer. Frontiers in Physiology. 2014;5:130.

[29] Moasser MM. The oncogene HER2: its signaling and transforming functions and its role in human cancer pathogenesis. Oncogene. 2007;26(45):6469–6487.

[30] Gilis J, Taelman S, Davey L, Martens L, Clement L. Pitfalls in re-analysis of observational omics studies: a post-mortem of the human pathology atlas. BioRxiv. 2020; 2020.03.16.994038. doi:https://doi.org/10.1101/2020.03.16.994038.

