## Supplementary Material for "Survival analysis of pathway activity as a prognostic determinant in breast cancer"

April 20, 2021

### Note S1: A null model based on set permutation

In the main text, we tested our pathways against the null model that there is a random association between gene expression, and the estimated pathway activity on one hand, and the survival of patients on the other hand. We have shown that our method is well-calibrated under that null.

However, empirical $p$ values can also be calculated against the null hypothesis that the sets of genes are grouped in a pathway are just randomly selected. This is a view that other pathway analysis methods have taken before, e.g. Gene-set enrichment analysis [2, 1].

To test significance under this revised null model, pathway-activities and Cox regressions were evaluated for random gene-sets for all investigated sizes of pathways. These permutations were used to calculate empirical $p$ values. To do this for a pathway of size n, we randomly sample n genes 10 000 times and run our analysis on this collection each time, storing the $z$ values from each regression. This forms the null distribution to which we compare the results of the curated pathway's result again, to derive $p$ values.

We obtained 56 significant pathways at a 5% FDR, which is a contrast to the 1030 pathways found under our main null model, yet provides further confirmation of the significance of our top pathways.

### References

[1] Jelle J Goeman and Peter Bühlmann. Analyzing gene expression data in terms of gene sets: methodological issues. *Bioinformatics*, 23(8):980–987, 2007.

[2] Aravind Subramanian, Pablo Tamayo, Vamsi K Mootha, Sayan Mukherjee, Benjamin L Ebert, Michael A Gillette, Amanda Paulovich, Scott L Pomeroy, Todd R Golub, Eric S Lander, et al. Gene set enrichment analysis: a knowledge-based approach for interpreting genome-wide expression profiles. *Proceedings of the National Academy of Sciences*, 102(43):15545–15550, 2005.

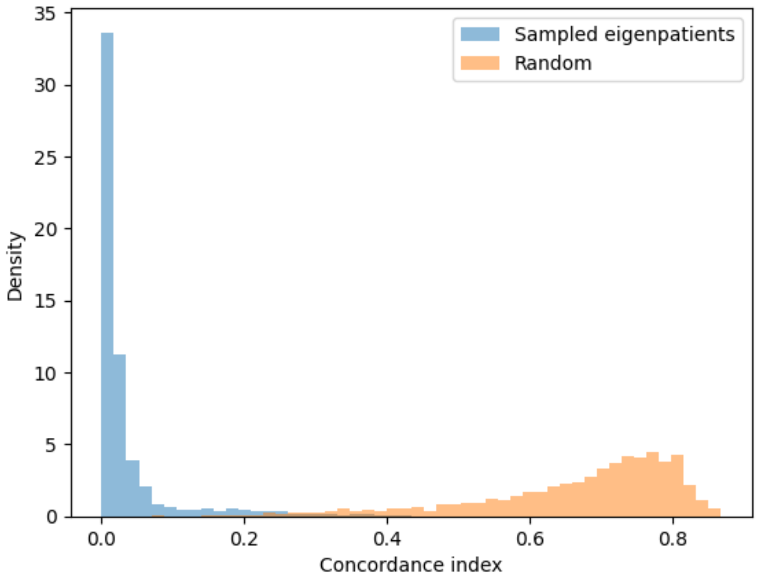


Figure S1: **Test of stability of pathways eigensamples.** We randomly selected a subset of 20% (398 samples) of the tumors and made pairwise comparisons of the direction of the eigensamples using the cosine distance. As a comparison, the procedure was repeated for randomly selected expression values picked from a uniform distribution, $\mathcal{U}(0, 1)$.

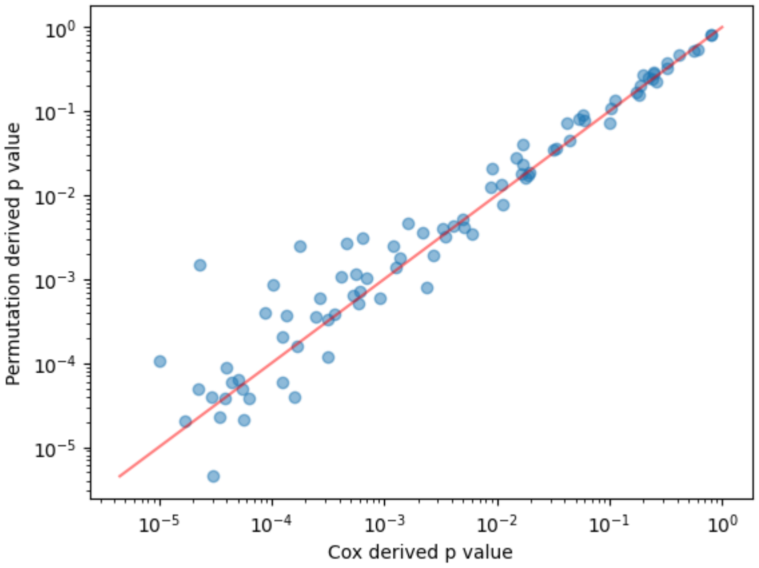


Figure S2: **Investigation of the statistical calibration of $p$ values from the Cox regression model.** The associations between gene expression values were permuted and the fraction of permutations with a more extreme outcome was compared to the Cox model's $p$ value.
